# The demographic decline of a sea lion population followed multi-decadal sea surface warming

**DOI:** 10.1101/2020.03.09.984716

**Authors:** Karen Adame, Fernando R. Elorriaga-Verplancken, Emilio Beier, Karina Acevedo-Whitehouse, Mario A. Pardo

## Abstract

**Background:** The population growth of top predators depends largely on environmental conditions suitable for aggregating sufficient and high-quality prey. We reconstructed numerically the population size dynamics of a resident population of California sea lions in the Gulf of California during 1978 – 2019, and its relation with the gulf’s multi-decadal sea surface temperature trend. This are the first long-term insights to the oceanic environment of the Gulf of California and to the population trend of one of its major predators.

**Results:** Our results indicate that a three-decade sustained warming explains statistically the population’s trend, including a ∼65% decline between 1991 – 2019, accounting for 92% of the variance. Long-term warming conditions started in the late 80’s, followed by the population’s decline, from a peak of 43,834 animals (range: 34,080 - 58,274) in 1991 to only 15,291 (range: range: 11,861 - 20,316) in 2019. The models suggested a century-scale optimum sea surface habitat for the population occurring in mildly temperate waters, from 0.18 to 0.39°C above the 100- year mean of 22.24°C.

**Conclusions:** The negative relation of the population size with warming sea surface conditions was evident, and the predictability of the former from the sea surface temperature 100-year anomalies was high. The mechanistic links of this relation are still untested, but an apparent diversification of pelagic fish catches suggests a reduction of high quality prey. We propose that this sea lion population should be considered as vulnerable to any disturbance that could add to the negative effects of the current sea surface warming conditions in the Gulf of California.

## Background

The physical structure of oceanic habitats often determines, through a multi-step process, the success of animal populations (Drinkwater et al., 2010). Sea surface temperature and its variability is widely known to affect top predators such as pinnipeds through bottom-up mechanisms (Melin et al., 2012; Moore, 2008; Trillmich and Ono, 1991). Abrupt or sustained changes of temperature affect the abundance and diversity of plankton communities (Bronikowski and Promislow, 2005), pelagic fishes (Chavez, 2003), and marine mammals (Laake et al., 2018; Simmonds and Isaac, 2007). The latter typically respond with alterations in foraging habits (Elorriaga-Verplancken et al., 2016; McClatchie et al., 2016), key physiological processes (Banuet-Martínez et al., 2017; Flores-Morán et al., 2017), reproductive success, and survival (Melin et al., 2012). Although many marine mammal populations can withstand and recover from short-term sea surface warming conditions (e.g. El Niño) (Melin et al., 2008), sustained positive environmental trends at a multi-decadal scale typically suppose shifts in the base of marine ecosystems (Anderson and Piatt, 1999), and arguably the diversity of potential prey. These shifts could lead to the decline of some marine mammal populations as the conditions move away from the optimum habitat to which the species have adapted (DeLong et al., 2017).

In the Gulf of California (also “the gulf” from herein), there is a resident population of California sea lions (*Zalophus californianus*) (Le Boeuf et al., 1983) genetically isolated from the other populations of the Northeast Pacific Ocean (Schramm et al., 2009). Although a ∼20% reduction of this population during the 1990’s was already proposed based on partial counts spanning 1997-2004 (Aurioles-Gamboa and Zavala-González, 1994; Szteren et al., 2006), its total size and temporal trend, as well as the historical environmental context, remain unknown. A recent review mentioned that the population decreased 44% between 1979 and 2016, based on unpublished data (Masper et al., 2019). Negative interaction with fisheries, and a temporal decline of the Pacific sardine (*Sardinops sagax*) in the early 1990’s, as a result of unspecified environmental changes, have been proposed as potential causes of the apparent decline (Szteren et al., 2006). Nevertheless, interannual events such as El Niño do not have a consistent correlation with the dynamics or health indicators of the gulf’s population (Aurioles and Le Boeuf, 1991; Hernández-Camacho et al., 2008). Given the uncertainty on both the apparent population decline and its drivers, we hypothesized that a long-term sea surface warming in the Gulf of California could be related, since this variable plays an important role in the thermodynamics of the entire basin and therefore would have the potential to trigger a progressive change in the base of the marine ecosystem, arguably altering the composition and diversity of prey for top predators.

To find support for this hypothesis, we made a numerical reconstruction of the total population size and its changes during the last 42 years (1978 - 2019), using individual trends of the 13 California sea lion reproductive colonies within the Gulf of California (Fig. 1), inferred from all available animal counts. Then, we explored the multi-decadal variability of the gulf’s sea surface temperature during the last 100 years (1920 - 2019), from which we were able to predict successfully the population’s multi-decadal dynamics. Nevertheless, it is important to point out that the lack of information did not allow us to test for the mechanisms involved in such habitat-species relationship, and therefore we can only propose hypothetical ecological scenarios to explain it based on similar responses already described for interannual or multiannual warm conditions.

**Figure 1.**
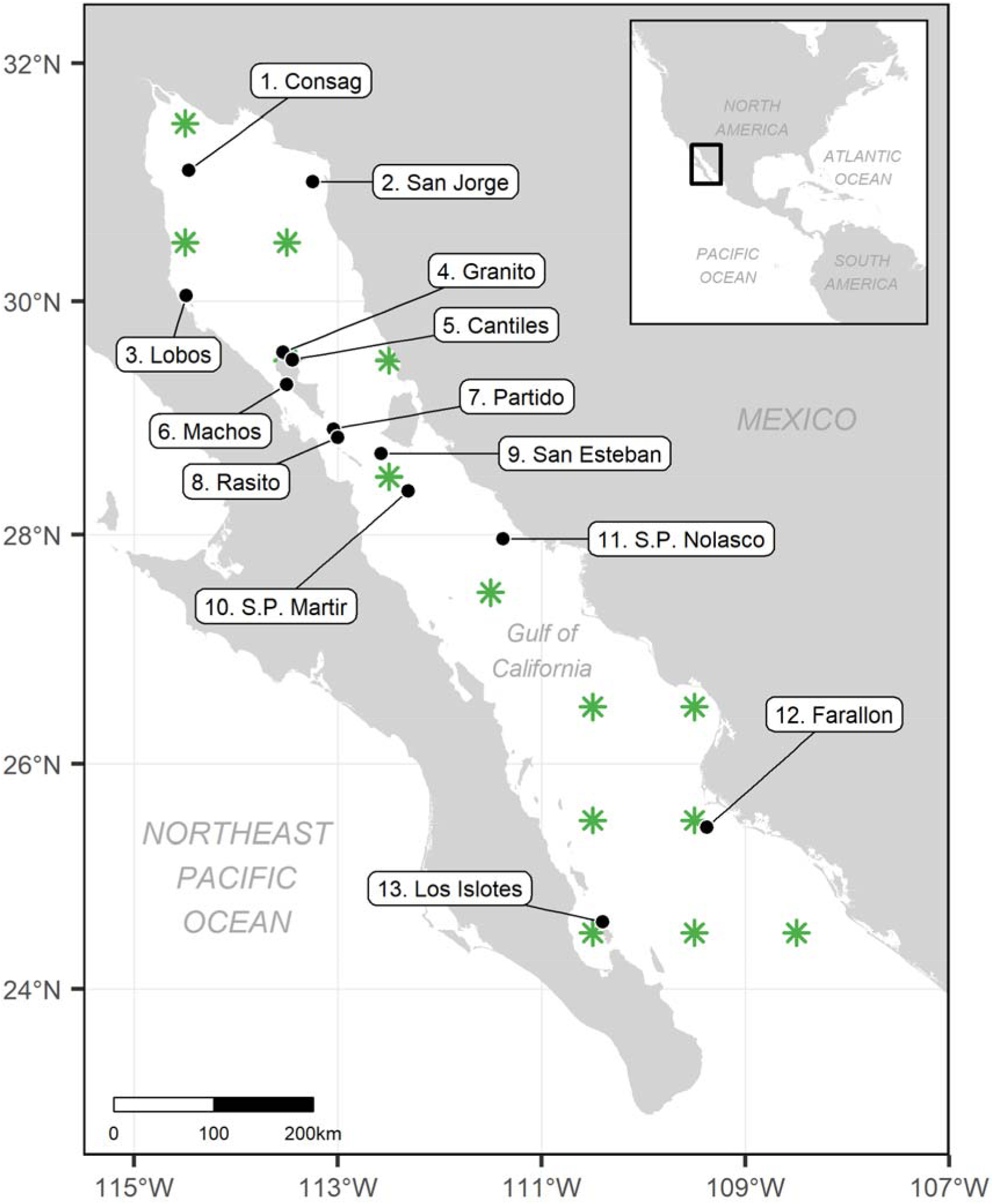
The Gulf of California in the Northeast Pacific Ocean. Black dots are the locations of the 13 reproductive colonies of California sea lions (*Zalophus californianus*), numbered from north to south. The green asterisks are the locations of the predicted sea surface temperature data used in this study.

The reproductive colonies in the Gulf of California are, from north to south: Rocas Consag, San Jorge, Lobos, Granito, Los Cantiles, Los Machos, El Partido, El Rasito, San Esteban, San Pedro Martir, San Pedro Nolasco, Farallon de San Ignacio, and Los Islotes (Le Boeuf et al., 1983; Szteren and Aurioles-Gamboa, 2011). Even though they are often grouped into four ecological regions (Szteren and Aurioles-Gamboa, 2011; Ward et al., 2010), or three genetic groups (Schramm et al., 2009), they function as individual units for the most part, because California sea lion individuals commonly exhibit natal philopatry to their colonies (Hernández-Camacho et al., 2008), which also reflects distinctive foraging habits (Porras-Peters et al., 2008). Our results confirmed a dramatic population decline in the last 28 years (1991 - 2019), after reaching its maximum in the early 1990’s. The gulf’s sea surface temperature anomalies from the 100-year mean (1920 - 2019) were able to predict the population size dynamics at a multi-decadal level and suggested an optimum habitat at mildly temperate conditions.

## Results

For all the 13 reproductive colonies, a second-order polynomial regression was the best model describing the abundance as a function of the year at a multi-decadal scale. The mean drone-based perception bias correction factor added 41% (range: 33 - 48) to the original boat-based counts. The mean percentage of non-pups in the population was 80% (range: 77 - 82), to which a 40.5% (range: 4.3 - 93.2) was added to account for animals likely at sea (Table 1). Almost all reproductive colonies exhibited a decreasing trend during the last three decades (1990’s - 2010’s), except for the southernmost small colony Los Islotes, which showed a sustained population growth in that period, with an apparent stabilization in the last four years (2015 - 2019) (Fig. 2). Some colonies showed decline even since 1978, but most of them, including the largest ones, showed a bell-shaped trend, with an increase during the 1980’s followed by a sustained decrease since the early 1990’s.

**Table 1.** Statistical summary of the posterior distributions of the most relevant parameters estimated by the models. The names in JAGS language are those of the code in Methods. A Gelman-Rubin statistic 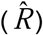 close to 1 indicates good convergence of chains. A high percentage of the effective number of iterations (*N*_*eff*_) indicates less uncertainty in the parameter’s estimation.

| Parameter | In JAGS code | Mean | SD | 2.5% | 25% | 50% | 75% | 97.5% | $\hat{R}$ | $N_{eff}$ (%) |
| --- | --- | --- | --- | --- | --- | --- | --- | --- | --- | --- |
| **Numerical reconstruction of the population size** |  |  |  |  |  |  |  |  |  |  |
| Population size in 1978 (1st of the series) | pred_sum_gulf[1] | 37651 | 5271 | 28728 | 33811 | 37139 | 41112 | 48755 | 1.001 | 100 |
| Population size in 1991 (highest peak) | pred_sum_gulf[14] | 45970 | 6428 | 35068 | 41281 | 45333 | 50192 | 59518 | 1.001 | 100 |
| Population size in 2019 | pred_sum_gulf[42] | 16006 | 2245 | 12217 | 14371 | 15787 | 17480 | 20736 | 1.001 | 100 |
| 1980's increase (%) | diff_decade_perc[1] | 15.570 | 0.918 | 13.784 | 14.947 | 15.571 | 16.190 | 17.373 | 1.001 | 83 |
| 1990's decrease (%) | diff_decade_perc[2] | -10.334 | 0.288 | -10.896 | -10.528 | -10.335 | -10.140 | -9.769 | 1.001 | 100 |
| 2000's decrease (%) | diff_decade_perc[3] | -31.994 | 0.378 | -32.736 | -32.247 | -31.995 | -31.738 | -31.252 | 1.001 | 100 |
| 2010's decrease (%) | diff_decade_perc[4] | -39.711 | 0.500 | -40.685 | -40.051 | -39.713 | -39.375 | -38.725 | 1.001 | 100 |
| 28-year decrease (%) (1991 - 2019) | diff_max_min | -65.179 | 0.522 | -66.196 | -65.532 | -65.183 | -64.828 | -64.149 | 1.001 | 100 |
| Proportion of undetected animals | prop_non_det | 0.444 | 0.004 | 0.436 | 0.441 | 0.444 | 0.446 | 0.451 | 1.001 | 35 |
| Proportion of non-pup animals | mu_prop_non_pups | 0.798 | 0.010 | 0.777 | 0.791 | 0.798 | 0.805 | 0.818 | 1.001 | 73 |
| Proportion of non-pups at sea | mu_at_sea_prop | 0.440 | 0.236 | 0.039 | 0.268 | 0.417 | 0.595 | 0.939 | 1.001 | 92 |
| Proportion at non-reproductive colonies | prop_other_rook | 0.060 | 0.002 | 0.057 | 0.059 | 0.060 | 0.061 | 0.063 | 1.001 | 72 |
| **100-year analysis of sea surface temperature ($SST_{ice}$)** |  |  |  |  |  |  |  |  |  |  |
| Mean 100-year $SST_{ice}$ | centu_mean | 22.241 | 0.109 | 22.026 | 22.167 | 22.241 | 22.314 | 22.455 | 1.001 | 100 |
| Mean 100-year $SST_{ice}$ anomaly | mu_sst_anom | 0.497 | 0.017 | 0.464 | 0.485 | 0.497 | 0.508 | 0.530 | 1.001 | 100 |
| **Ecological model (population size response to $SST_{ice}$ anomalies)** |  |  |  |  |  |  |  |  |  |  |
| Intercept | a0 | 3.773 | 0.034 | 3.705 | 3.751 | 3.774 | 3.796 | 3.838 | 1.001 | 72 |
| First coefficient | a1 | 35.987 | 48.552 | 0.846 | 3.168 | 12.379 | 48.897 | 174.368 | 1.006 | 0.38 |
| Second coefficient | a2 | -1.367 | 0.178 | -1.714 | -1.487 | -1.368 | -1.247 | -1.016 | 1.001 | 17 |
| Mixing parameter | theta | 0.824 | 0.236 | 0.137 | 0.756 | 0.938 | 0.984 | 0.996 | 1.004 | 0.72 |
| Bayesian R-squared | r_squ_ecol | 0.915 | 0.012 | 0.881 | 0.910 | 0.918 | 0.923 | 0.927 | 1.001 | 100 |

**Figure 2.**
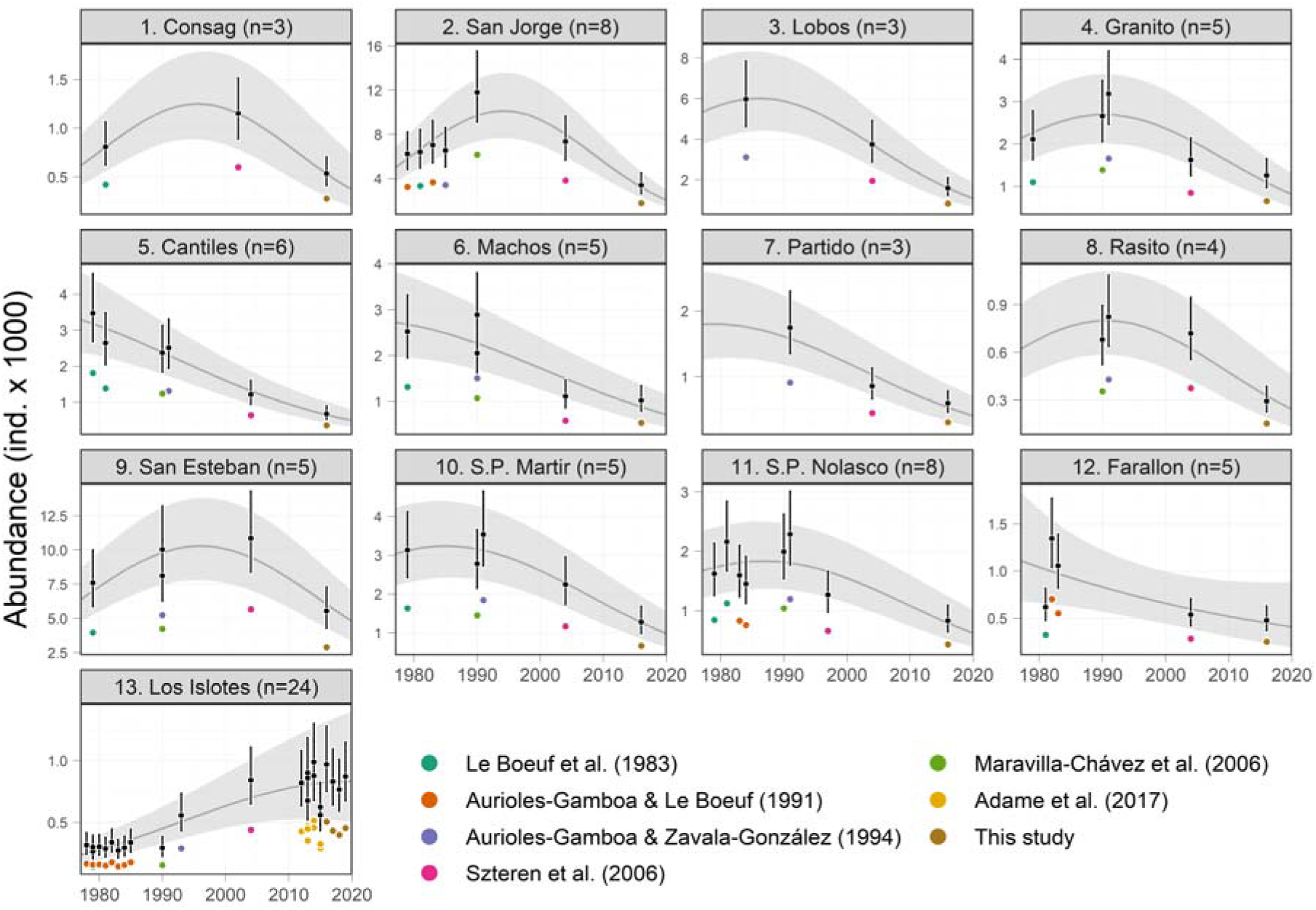
Abundance trends of California sea lions (*Zalophus californianus*) in the Gulf of California at the 13 reproductive colonies. Colored dots represent the original boat-based counts with the reference that reported them. Black dots and error bars represent the medians and the 95%-credible intervals (CIs) of the estimated abundance, accounting for perception and availability biases. Medians and 95%-CIs of the predicted trend are shown as dark gray lines and shaded areas, respectively. The colonies are numbered from north to south as portrayed in Fig. 1, along their sample sizes.

The most populated colonies predicted for 2019 were San Esteban with 4,959 animals (range: 3,308 - 7,633), San Jorge with 2,236 animals (range: 1,476 - 3,476), and Lobos with 1,175 (range: 677 - 2,100). The smallest were El Rasito, Partido, and Consag, with abundances of less than 500 animals. The sum of the colonies’ annual predictions was augmented by 6% (range: 5.7 - 6.3), corresponding to the estimated proportion of animals at the 16 non-reproductive colonies during a typical reproductive season (Table 1). According to this numerical reconstruction, the population had 35,972 animals (range: 27,892 - 47,682) in 1978. During the 1980’s, it gained 15.5% (range: 13.7 - 17.4), reaching its historical maximum in 1991 with 43,834 animals (range: 34,080 - 58,274). From that peak, the population lost 10.3% (range: 9.8 - 10.9) during the 1990’s, 32% (range: 31.3 - 32.7) in the 2000’s, and a 39.7% (range: 38.7 - 40.7) in the 2010’s. For 2019, the population size would have been of 15,291 animals (range: 11,861 - 20,316), which represents a 65.2% decline (range: 64.1 - 66.2) in the last 28 years (1991 - 2019) (Fig. 3; Table 1).

**Figure 3.**
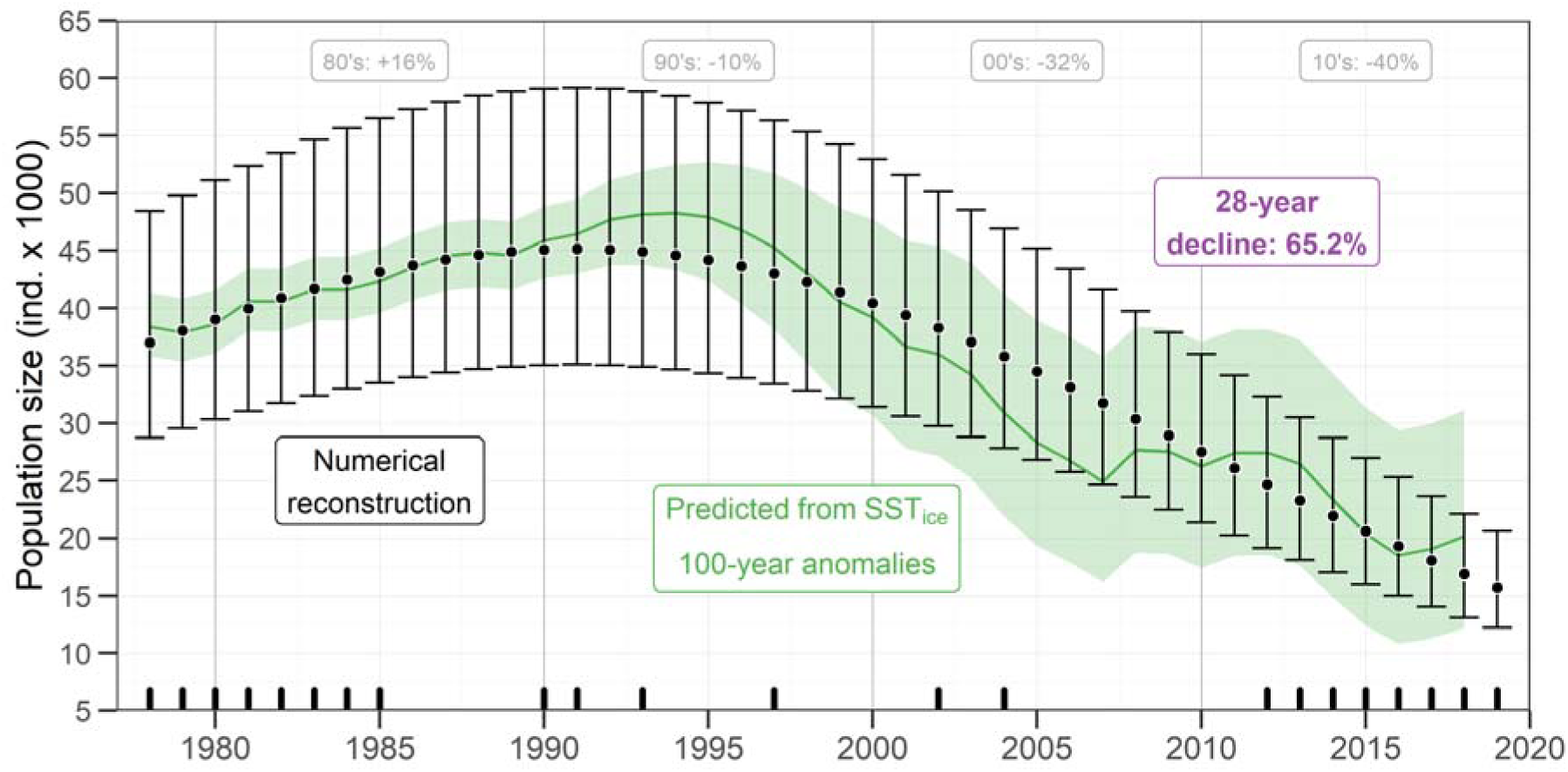
Historical population size of California sea lions (*Zalophus californianus*) in the Gulf of California. Annual medians (black dots) and 95%-credible intervals (CI; black error bars) of the numerical reconstruction are shown. Black inner ticks on the X axis mark the years for which any colony count was available. Medians of decadal changes and the 28-year decline in percentage are within gray and purple boxes, respectively. The population size prediction from the multi-decadal anomalies of sea surface temperature (SST_ice_) is shown as a green thick line (median) and green shaded area (95%-CI).

The time series of SST_ice_ anomalies from the 100-year mean of 22.2 °C (range: 22.02 - 22.45 °C) showed a warming of the Gulf of California since the late 1980’s (Fig. 4). It was especially steep during the 1990’s, and less accelerated during the 2000’s and the 2010’s, reaching +1.06°C in 2019 (range: +0.85 - +1.28). The minimum temperatures occurred during the mid-1930’s, reaching anomalies of −0.57°C (range: −0.77 - −0.35). During 1991, when the California sea lion abundance reached its maximum, the median SST_ice_ anomaly was +0.11°C (range: −0.11 - +0.31) (Fig. 4).

**Figure 4.**
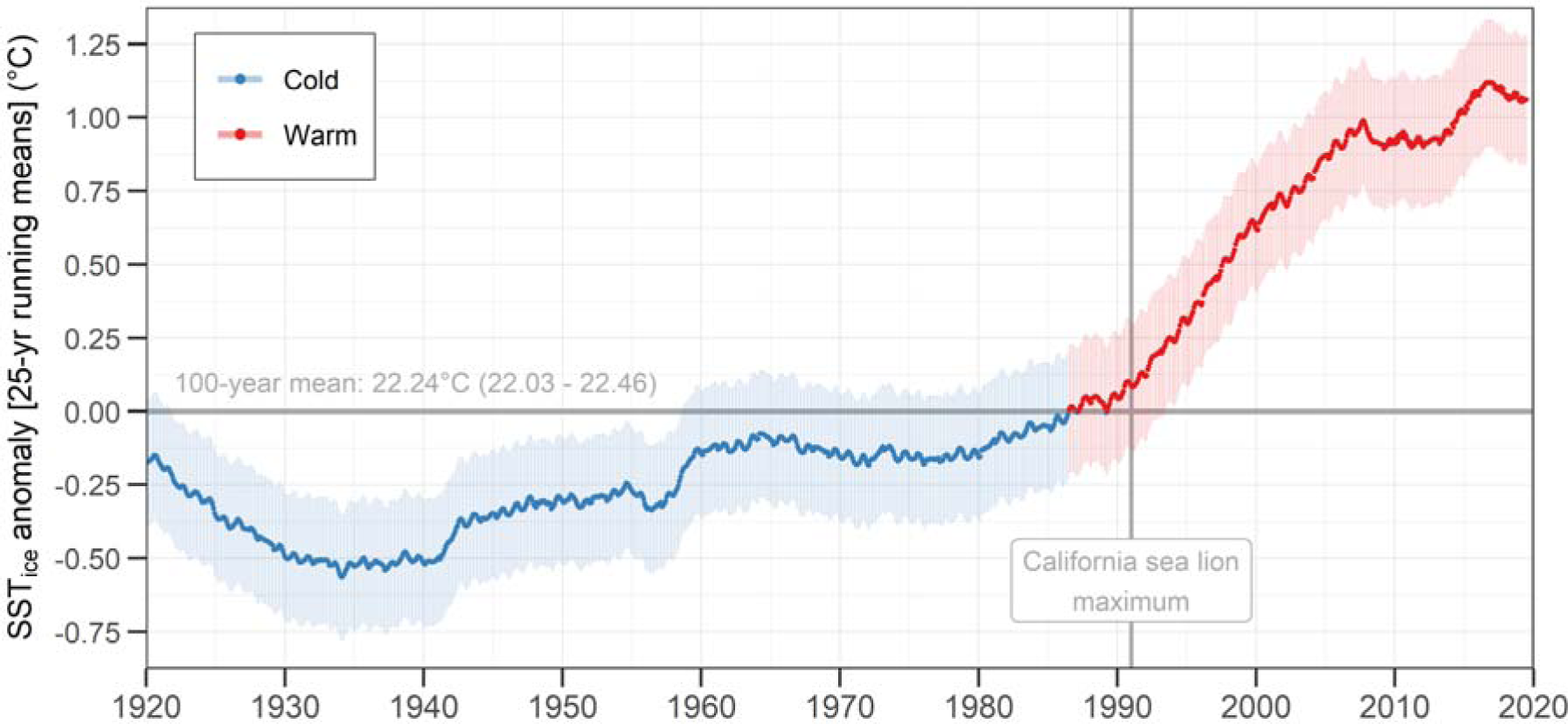
Sea surface temperature (SST_ice_) anomalies from the 100-year mean (horizontal gray line) in the Gulf of California. The colored thick line represents the median and the shaded area is the 95%-credible interval. The locations of SST_ice_ estimations are portrayed in Fig. 1. The maximum California sea lion (*Zalophus californianus*) population size occurred in 1991 (Fig. 3).

According to our model, the population size of California sea lions in the Gulf of California can be predicted successfully from the SST_ice_ 100-year anomalies, at a multi-decadal scale (i.e. 25-year running means). After testing for several degrees of curve complexity, the best model describing this relationship was a mixture between a second-order polynomial and a parabola function. This was accomplished through a mixing parameter, whose median closer to 1 (0.77; range: 0.06 – 0.99) indicated that the parabola function contributed with most of the fit (Table 1). The proportion of the variance explained by this model (i.e. the Bayesian R-squared) was 0.918 (range: 0.880 - 0.926), suggesting an extremely high predictability. The shape of the resulting curve (Fig. 5) implies that the population size is expected to decrease at both very high and very low SST_ice_ anomalies, relative to the 100-year mean. According to the predictions of this model (median and 95%-CI), the range at which the population size would reach its maximum would be during anomalies from +0.17 to +0.40°C (i.e. 25-year running means of SST_ice_ between 22.49 and 22.72°C) (Fig. 4). This could be interpreted as the population’s optimum physical habitat conditions. Additionally, the population size would have reached half of its maximum bellow ∼23°C, or a maximum anomaly of +0.76°C from the 100-year mean.

**Figure 5.**
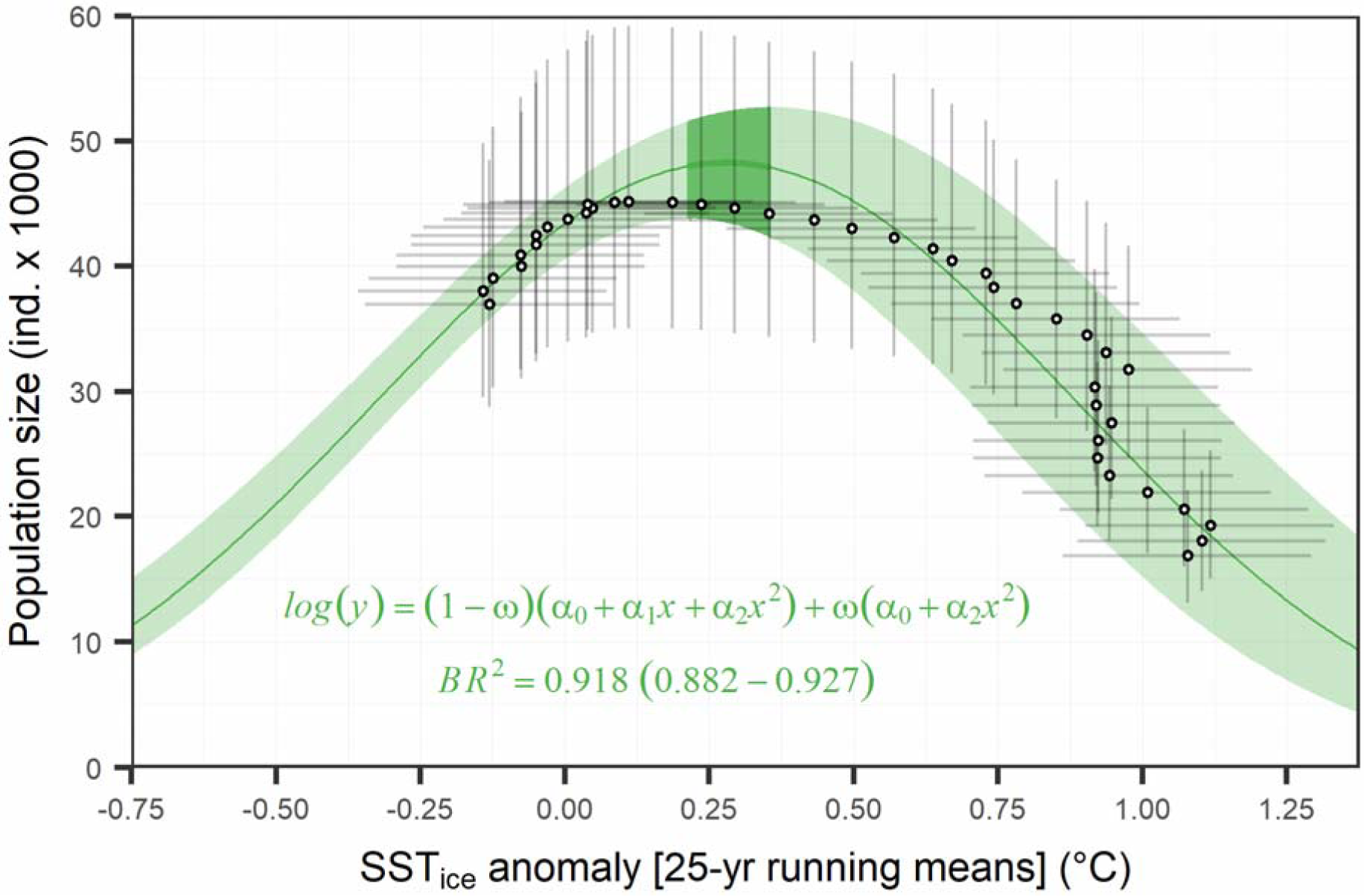
The California sea lion (*Zalophus californianus*) population size in the Gulf of California as a function of sea surface temperature (SST_ice_) multi-decadal anomalies from the 100-year mean of 22.24°C (range: 22.02 – 22.45). The green line and shaded area are the median prediction and the 95%-credible interval (CI), respectively. The darker line and shaded area represent the multi-decadal habitat optimum, between +0.17 and +0.40°C. Black dots and thin error bars are the medians and 95%-CIs of the annual estimations of both variables. The posterior summaries of the equation’s parameters are portrayed in Table 1.

The predictions of the annual population size using the posterior coefficients (Table 1) of the habitat model confirmed its high predictability. Though with higher uncertainty for the last two decades (i.e. during warmer conditions), the model predicted successfully the median population sizes that resulted from the numerical reconstruction (Fig. 3). Moreover, since this ecological model was based on SST_ice_ data at a higher temporal resolution than that of the animal counts, it was also capable of showing some details in the expected trend that the numerical reconstruction could not. The most important difference was that the numerical reconstruction tended steadily towards lower values, whereas the abundances predicted from the SST_ice_ anomalies suggested some level of stabilization of the trend during the late 2000’s and the 2010’s, or at least, a less pronounced decrease, compared to that of the 1990’s and early 2000’s (Fig. 3).

## Discussion

The negative statistical relation of the multi-decadal SST_ice_ warming in the Gulf of California (1990’s - 2010’s) with the California sea lion population trend was evident. This environmental condition predicted accurately the annual population size and trend, accounting for 91.8% of the variability (range: 0.882 - 0.927). California sea lions reflect long-term changes in the marine environment (Le Boeuf et al., 2002; Melin et al., 2012) because, as marine mammals and top predators in general, they are conspicuous, with high energetic demands, and have a long lifespan (Moore, 2008). Therefore, it is possible that other predators within the gulf have also reacted to this scale of environmental variability. Such effects at the multi-decadal scale cannot be interpreted as equivalent to those that have been observed in response to inter-annual warm events such as El Niño, which are well described (Aurioles and Le Boeuf, 1991; Gerber and Hilborn, 2001; Ono et al., 1987), and typically include a strong generalized depletion of prey availability and drastic changes in foraging habits (DeLong and Antonelis, 1991; Elorriaga-Verplancken et al., 2016; Trillmich and Limberger, 1985). Compared to these short-term shifts, the knowledge of the effects of long-term environmental changes is very limited (Fritz and Hinckley, 2005; Merrick et al., 1987; Trites and Donnelly, 2003), yet it is essential to understand and predict population dynamics under climate change scenarios.

While the negative effects of El Niño events tend to last for only two or three years (Aurioles and Le Boeuf, 1991; Trillmich and Limberger, 1985), a multi-decadal warming would suppose a progressive, sustained, habitat change affecting multiple generations. It is challenging to understand the mechanisms involved in the response of a predator population to this scale of sea surface warming, to the point of reducing dramatically its size. One of the possible causes of a long-term decline in a spatially restrained pinniped population could be a diet change from high quality prey to what has been referred to as “junk food” (Fritz and Hinckley, 2005; McClatchie et al., 2016). When preferred prey becomes less available, predators must resort to a less nutritious diet. In the California Current Large Marine Ecosystem, California sea lions feed commonly on small pelagic fishes, including several species of sardine, anchovy, and mackerel (Sweeney and Harvey, 2011), which are considered of high quality because of their high calorie and fat contents, in comparison to other prey such as the market squid (*Doryteuthis opalescens*) or the rockfish (*Sebastes jordani*), which are more common prey items during warming conditions (McClatchie et al., 2016; Sweeney and Harvey, 2011). In that region, this type of prey replacement has resulted in pups with lower body masses.

An analogous prey replacement could have happened in the Gulf of California in 1978 - 2019, although this California sea lion population has different foraging habits than those of the California Current Large Marine Ecosystem, and few studies have described its diet (Garcia-Rodriguez and Aurioles-Gamboa, 2004; Porras-Peters et al., 2008). The latter varies between colonies and consists on a large variety of species, including high-trophic level items such as the midshipman (*Porichthys* sp.) and squid (*Leachia* sp.), as well as low-trophic level species like the Pacific jack mackerel (*Trachurus symmetricus*), the Pacific sardine (*Sardinops sagax*), and the northern anchovy (*Engraulis mordax*). These prey were found in scats collected in the mid 1990’s and early 2000’s, so the composition of the diet in the late 1980’s, before the population decline, is unknown. However, some trends in the fishery of small pelagic species could support the idea of a shift in the base of the trophic web. Sardine fisheries in the Gulf of California increased slowly until 1989; then, a massive collapse occurred. The catches plummeted from almost 300,000 tons to 7,000 tons in less than three years. Since then, several rises and falls have occurred, including the most extreme in 2008, when catches fell from over 500,000 tons to 3,000 tons in five years (Velarde et al., 2015). Nevertheless, as sardine fisheries plummeted and their trend became more unpredictable, other small pelagic fishes such as the thread herring (*Opisthonema libertate*), the Pacific mackerel (*Scomber japonicus*), the northern anchovy, and the bigmouth sardine (*Cetengraulis mysticetus*), gained relative importance in the catches (Lanz et al., 2009). This could indicate a diversification of the potential prey for California sea lions as the sea surface temperature raised. The effect of ocean surface warming on global fisheries is well documented, particularly in temperate regions, where warmer water species are being captured at higher latitudes (Cheung et al., 2013; Vergés et al., 2016). Nevertheless, since the coverage of available data of potential prey is not comparable to the multi-decadal scale of this study, nor is spatially representative of the entire gulf, and given the high heterogeneity of fishing effort, it is not possible to validate this hypothetical ecological mechanism.

If California sea lions have been foraging on more diverse but lower quality prey in the Gulf of California for several generations, it is expected that the main drivers of the population dynamics, fertility and/or survival rates, would have lowered. Although there is no direct evidence of this, the mean proportion of pups during boat-based counts before the decline (1979 - 1991) was ∼25%, and ∼42% for adult females (Aurioles-Gamboa and Zavala-González, 1994). In 2016, we found a proportion of pups of only 15% whereas the proportion of adult females was similar (∼42%) (Supplementary Information, Table S1), suggesting a reduction in fertility or pup survival that should be further explored. In contrast, in a colony of the species in the California Current Large Marine Ecosystem, with a positive population trend until 2014, the proportions of pups and females were ∼41% and ∼40%, respectively (Elorriaga-Verplancken et al., 2015). Fertility and pup survival reductions are common consequences of short-term warm anomalies (McClatchie et al., 2016; Melin et al., 2008; Trites and Donnelly, 2003), but their link to multi-decadal environmental shifts is unknown. An increased foraging effort by juvenile and adult animals in warmer conditions should not be discarded as a complementary factor that could explain the decline of California sea lions on land. During warm anomalies, telemetry studies have shown that this species displays offshore foraging trips up to three times longer, presumably as a consequence of reduced prey availability (Costa et al., 2004; Weise et al., 2006), resulting in fewer animals on the islands when counts are made. Further telemetry data are needed to build a dynamic correction factor of animals likely at sea.

The 15.5% increase (range: 13.7 - 17.4) of the population size during the 1980’s (Fig. 5) was also successfully predicted by SST_ice_ 100-year anomalies. Because the shape of the curve of that model was unimodal, the population size would also be lower under extreme cold anomalies, at the multi-decadal scale. The optimum sea surface temperatures revealed by this model could hypothetically benefit the aggregation of high quality prey in the Gulf of California, such as sardine, whose fishery also increased during the 1970’s and early 1980’s (Lluch-Belda et al., 1986; Nevárez-Martínez et al., 2001; Velarde et al., 2015). Though it could be argued that this California sea lion population increased during that period as a result of its recovery from documented legal hunting prior to the 1970’s (Cass, 1985; Masper et al., 2019; Zavala-González and Mellink, 2000), there is no evidence that the level of hunting was high enough to produce an appreciable decline of the population from which it would have recovered, at least not enough for a bottleneck effect on the population (Gonzalez-Suarez, 2010). Moreover, if that were the case, the SST_ice_-based model would not be able to predict such hypothetical recovery.

One particular finding of this study, and the exception to the general pattern, was the abundance trend at the colony Los Islotes, which was the only one that showed a steady increase during the study period, tending to stabilize in the last five years. It is the southernmost reproductive colony of the Gulf of California, located in La Paz Bay, a region characterized for its high biological production year round, attributed to a local mesoscale gyre phenomenon occurring during summer (Martínez-López et al., 2001; Pardo et al., 2013). This means that this colony would benefit from a more stable availability of prey. Also, the colony’s prey has historically been reported as be of higher trophic levels compared to others (Porras-Peters et al., 2008). As a result, the impact of hypothetical variations of small pelagic fish may be lower on this particular region.

## Conclusions

More studies focused on the foraging habits and reproductive health of the population, as well as studies on the abundance and diversity of potential prey, are needed to reach satisfactory conclusions on the mechanisms driving the population’s long-term dynamics. It is also probable that other predators in the Gulf of California that depend on similar low-trophic-level prey would show similar population trends, but to our knowledge, this is the first study quantifying such temporal scale of environmental variability and biological response in the gulf. Based on our results, we propose that the California sea lion population of the Gulf of California should be considered as vulnerable to any disturbance that could add to the apparent negative effects of the current sea surface warming conditions. Such conservation status should be maintained until the population size reaches at least half of the maximum predicted during the early 1990’s, which our models predict would occur at a 25-year running mean SST_ice_ bellow ∼23°C, that is, a maximum anomaly of +0.76°C from the 100-year mean.

## Methods

All the analyses described below were based on algebraically explicit Bayesian regression models (Kéry and Royle, 2016; McCarthy, 2007), whose parameters of interest were estimated as samples from their posterior distributions through a Markov Chain Monte Carlo procedure, implemented in the language Just Another Gibbs Sampler (JAGS) (Plummer, 2003), in R (R Core Team, 2018). This approach avoids the loss of information through the propagation of uncertainty between connected estimations and models, accounting effectively for scarce and/or sparse data, if needed. We ran one million iterations in five independent chains, retaining every 20th value to avoid autocorrelation, and discarding the first 20% as a burn-in phase for each chain (see detailed algebraic explanations and JAGS codes below). The results of the estimations are reported throughout the text as their medians, accompanied by ranges that represent the 95%-credible intervals of the posterior distributions. All the data needed to run the models is provided fully arranged as three R’s data list objects in Supplementary Information, File S1.

### Population abundance and trend

When estimating abundance of pinnipeds, pup counts are typically used to infer those of other age classes from life charts that take into account life expectancy and fertility rates, among other parameters (Kirkwood et al., 2005; Laake et al., 2018; Lowry et al., 2014). Unfortunately, only part of the available historical counts of California sea lions in the Gulf of California contains information on pups (Supplementary Information, Table S1), and there are no life charts specifically estimated for each reproductive colony (Masper et al., 2019); therefore, such traditional estimates were not feasible. Instead, we focused on total counts, i.e. all age/sex classes, as a consistent parameter that can be also used to estimate pinniped abundance (Aurioles-Gamboa and Zavala-González, 1994; Elorriaga-Verplancken et al., 2015; Szteren et al., 2006), especially if some perception and availability biases are accounted for (Adame et al., 2017; Bonnell and Ford, 1987).

The main challenge for estimating the population size of California sea lions and its multi-decadal trend in the Gulf of California was that there is only one reproductive season, in 2016, for which there are available counts of animals in all of the 13 reproductive colonies, an effort of this study for which we implemented both visual and drone-based counts. The rest of the available counts spanning the 42-year time series (1978 - 2019), both published and described here for the first time, were made separately for different colonies during different reproductive seasons, spanning 1978 - 2019. Therefore, it was impossible to estimate more than one total abundance based on the simple sum of counts from all colonies. Fortunately, although very sparse in time, these 86 counts followed the same boat-based protocols (Adame et al., 2017; Aurioles-Gamboa and Zavala-González, 1994; Le Boeuf et al., 1983; Szteren et al., 2006), and therefore were comparable. This method consists in circumnavigating the colonies during morning hours, at 15 - 45 m from the shoreline, with hand-held binoculars, at a speed of 5 - 7 km h^-1^. The counts included all animals detected on land, as well as those swimming near the surface between the boat and the colony.

To solve perception bias and correct categorization errors, 16 drone-based counts were made parallel to the boat-based counts during the 2016 reproductive season (Supplementary Information, Table S1), taking aerial photographs of all the areas with presence of California sea lions, at an altitude of ∼20 m and the best possible angle of view (see detailed protocol in Adame et al., 2017). The categorization of California sea lions followed established guidelines for identifying each individual as adult male, sub-adult male, adult female, juvenile, pup, or undetermined (Le Boeuf et al., 1983; Lluch-Belda, 1969; Peterson and Bartholomew, 1969). The counts reported for the first time in this study were made during 2016 - 2019 under research permits SGPA/DGVS/00050/16-19 issued by *Dirección General de Vida Silvestre* - *Secretaría de Medio Ambiente y Recursos Naturales*, with authorization of *Comisión Nacional de Áreas Naturales Protegidas*.

The first step of the analysis was fitting individual regressions of animal boat-based counts at each reproductive colony as functions of the year (1978 - 2019), only during breeding seasons. We tested for different degrees of curve complexity (i.e. incremental polynomial orders), choosing the best fit according to the lowest value of the Deviance Information Criterion (DIC) (Gelman et al., 2014). A Poisson likelihood was stated for all counts with a logarithmic link function (see model equations below). We used all published counts, and those reported for the first time in this study (Supplementary Information, Table S1). Given that most of the counts were very sparse in time during the 42-year study period (1978 – 2019), these curves were intended to identify multi-decadal variations only, rather than interannual or more fine-scale dynamics. However, the Bayesian framework of the analyses assured that the uncertainty associated to the estimations captures accurately the number of observations and their sparsity in time, which is especially the case of Consag, Lobos, and Partido.

The perception bias of boat-based counts was estimated for all available surveys as the proportion of animals detected from the boat with respect to those detected from the aerial drone photos at each reproductive colony. All proportions were stated as binomial likelihoods, whose inverse main parameters were used as additive correction factors for the annual abundance predictions. We also estimated an availability bias correction factor to add the proportion of animals likely to be at sea when the surveys were made. For that, we used a mean of those proportions reported in a study that compared on-land and at-sea aerial counts at the colonies of the same species in the California Current Large Marine Ecosystem (Bonnell and Ford, 1987). Unfortunately, there was no available information for our study area for this purpose. Since those proportions were with respect to all categories except pups, we had to estimate first the mean proportion of the former in the Gulf of California, from 76 counts with available information of pups (Supplementary Information, Table S1). Then, the correction factor was applied only to non-pup animals for each reproductive colony. Although the proportion of animals foraging at sea can vary as a function of prey availability (Kuhn and Costa, 2014; Weise et al., 2006), there was no available data or previous information that allowed us to address this dynamically. Therefore, our model assumed this proportion as constant. Since El Partido and El Rasito did not have enough observations during the first 10 years of the time series, and given that both colonies have very limited area available for California sea lions, we set upper truncation limits for their predictions at the beginning of the time series to their maximum total counts available.

To obtain the annual posterior distributions of the total population size, we summed the annual predictions of abundance at each reproductive colony from 1978 to 2019, all within the same hierarchical Bayesian structure to propagate the uncertainties of the colonies’ estimations (Kéry and Royle, 2016). We also added a final availability bias correction factor to account for the animals likely present at the 16 known non-reproductive colonies during a typical reproductive season. We estimated this proportion based on the counts reported by the only study that included such colonies (Aurioles-Gamboa and Zavala-González, 1994). To better visualize the population size dynamics, we also estimated decadal and maximum percentages of change during the 42-year time series.

### 100-year anomalies of sea surface temperature

We estimated 25-year running means of sea surface temperature within the Gulf of California and their anomalies from the 100-year mean, from 1920 to 2019 (n = 1193). This allowed us to filter interannual and decadal signals, such as El Niño Southern Oscillation (ENSO) or the Pacific Decadal Oscillation (PDO), respectively, keeping only multi-decadal variability, in accordance to the scale of the California sea lion population trend explored in this study. The dataset consisted of predictions from a two-stage reduced-space optimal interpolation procedure, and the superposition of quality-improved gridded sea surface temperature observations onto reconstructions, based on historical ice concentrations at the Earth’s poles (Rayner et al., 2003). To differentiate it from the traditional remotely-sensed sea surface temperature (SST), we refer it as SST_ice_. The product was developed by the Met Office Hadley Centre (http://hadobs.metoffice.com/) for use in climate monitoring and modeling, and is freely distributed at a monthly-one-degree resolution by the Environmental Research Division’s Data Access Program of the National Oceanic and Atmospheric Administration (https://coastwatch.pfeg.noaa.gov/erddap/griddap/erdHadISST.html), for which 14 SST_ice_ prediction points corresponded to the Gulf of California (Fig. 1).

### Habitat-based population trend

We fitted a regression model of the 42 annual predictions of California sea lion population size as a function of the SST_ice_ 100-year anomalies, at an annual basis, using the means and standard deviations of the posterior distributions of both variables estimated by the two models described above. We tested for incremental polynomial degrees for the regression, but also for pairwise combinations of models through a mixing parameter (Hoeting et al., 1999), which can add flexibility to the curve and increase the explained variability. The best fit was chosen based on the lowest DIC. The results of this function allowed us to calculate geometrically a range of optimum habitat, which was interpreted as those SST_ice_ anomalies values at which the population size predictions reached the maximum at the lower and upper 95%-credible intervals of the curve (Fig. 5). The same approximation was used to calculate the minimum value of the predictor at which the population size would reach half of its maximum.

### Algebraic and JAGS code details for all models

For the numerical reconstruction of the California sea lion population size in the Gulf of California, all colony counts (*C*) available (*i*) were assumed to come from a Poisson likelihood with a logarithmic link, whose parameter (*λ*) was a function of the year (*y*) at each reproductive colony (*j*), which had a fixed effect on the relation:

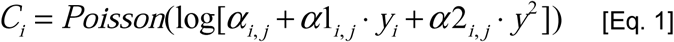

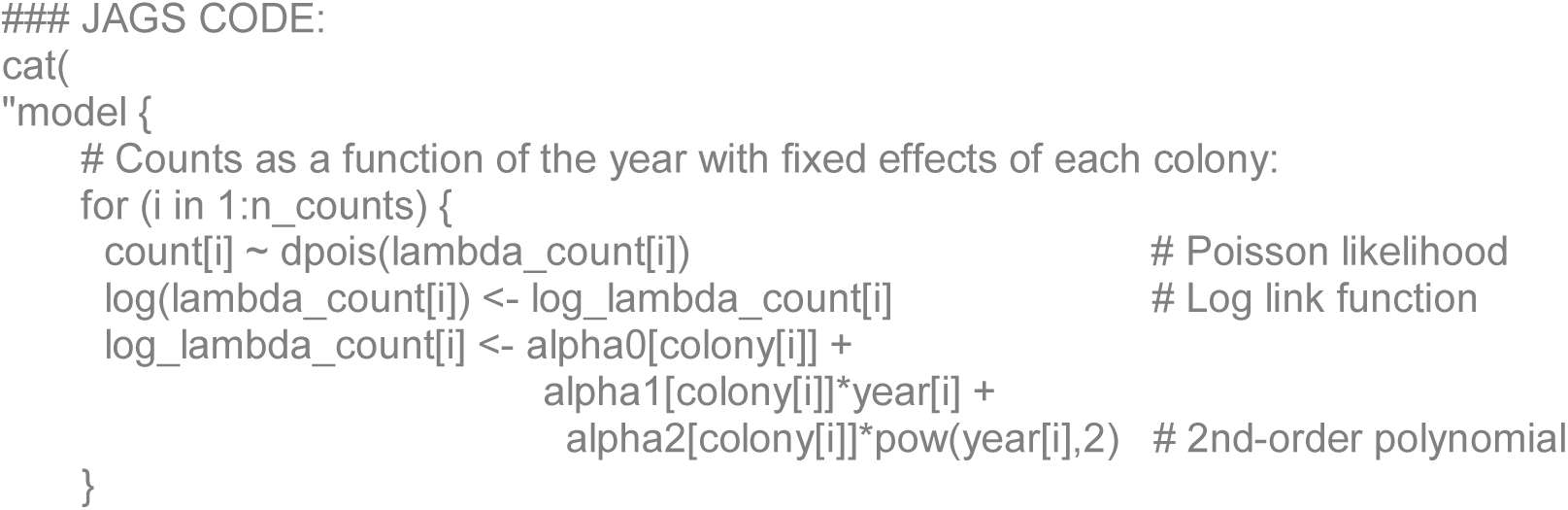

For each reproductive colony (*j*), boat-based visual counts (*V*) in 2016 came from a Binomial likelihood, with a mean proportion of detected animals (*P*_*det*_) respect to the total (*T*_*drone*_) (i.e. drone-based). The complement proportion of the former were the un-detected animals (*P*_*und*_):

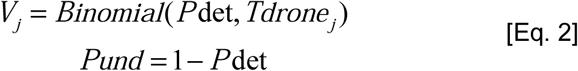

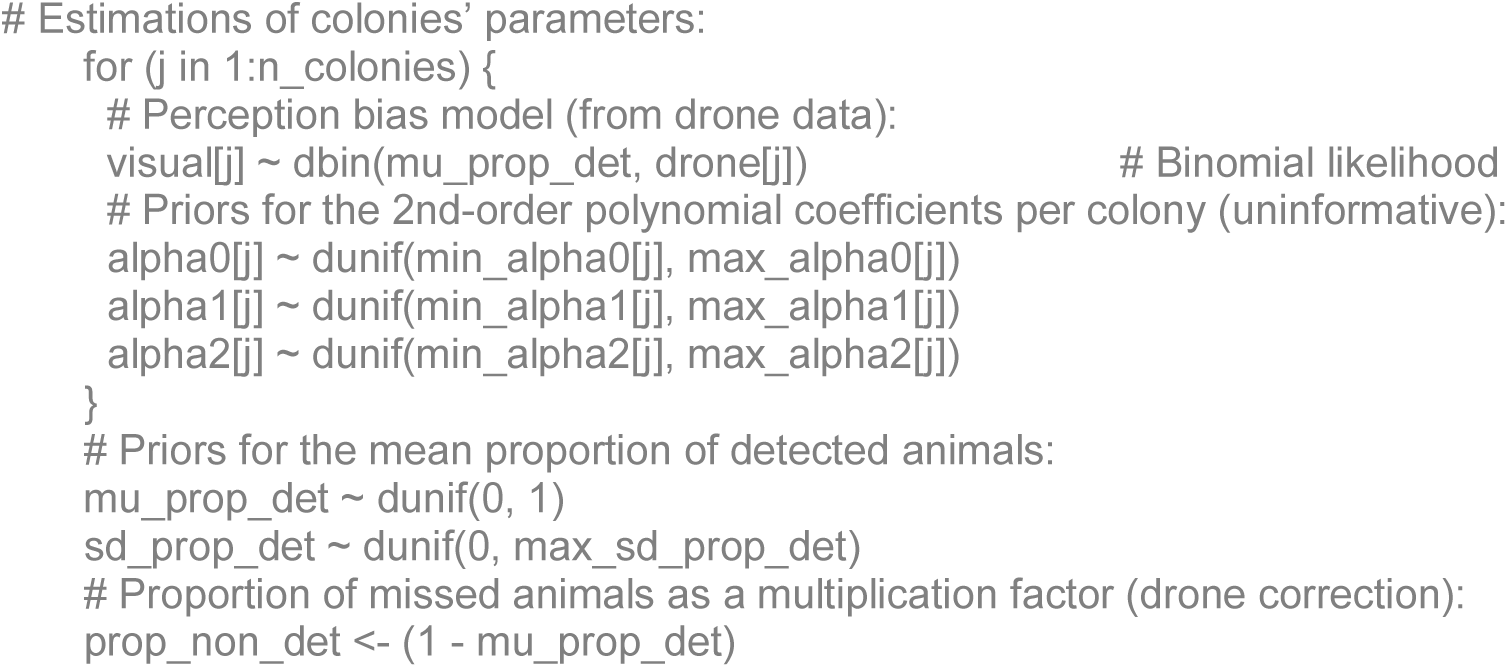

For each survey (*k*) with available information of pups, non-pup counts (*Cnp*) were stated to come from a Binomial likelihood with a mean proportion (*P*_*np*_) respect to the total (*T*):

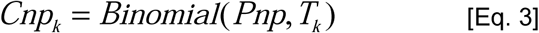

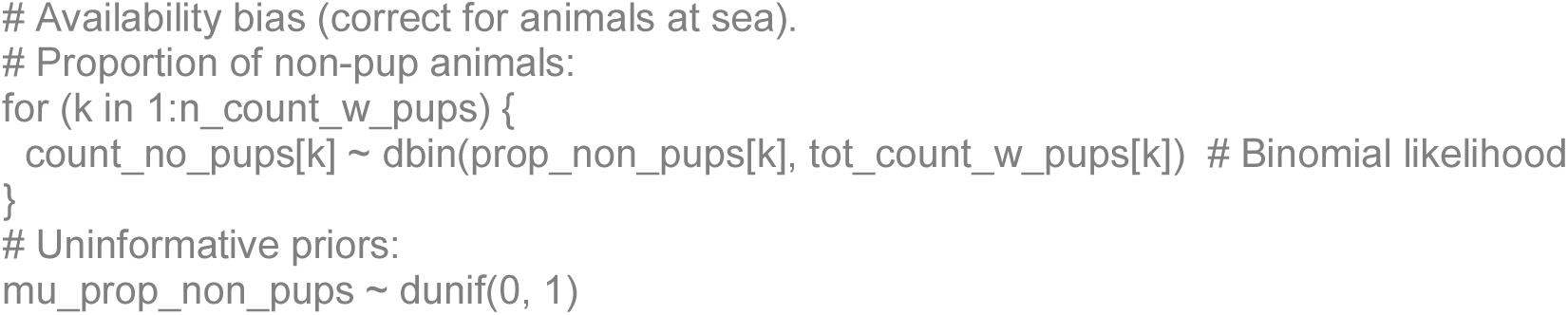

For each reference (*l*) of the proportions published of adult animals at sea (*P*_*ats*_), a normal likelihood was stated with unknown mean (*µ*_*ats*_) and standard deviation (*σ*_*ats*_):

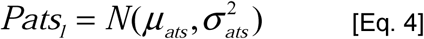

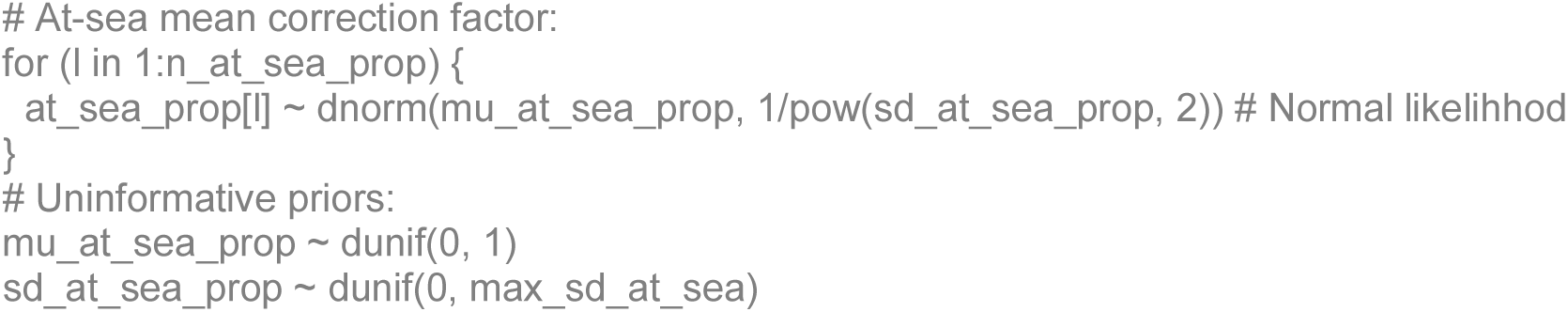

The only total count (*TC*) available encompassing all reproductive and non-reproductive colonies was stated as the second parameter of a binomial likelihood whose number of animals at the 16 non-reproductive colonies (*N*_*nrc*_) represents a mean proportion *P*_*nrc*_:

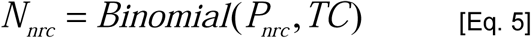

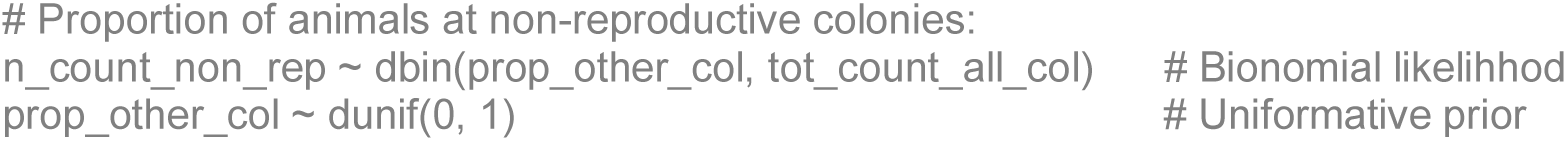

The annual (*y*) predictions of animal abundance (*A*) for the 13 reproductive colonies (*r*) summed were derived from the function in Eq. 1, adding the proportion of undetected animals (*P*_*und*_) in visual surveys (i.e. drone-based correction) (Eq. 2):

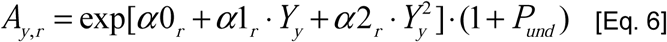

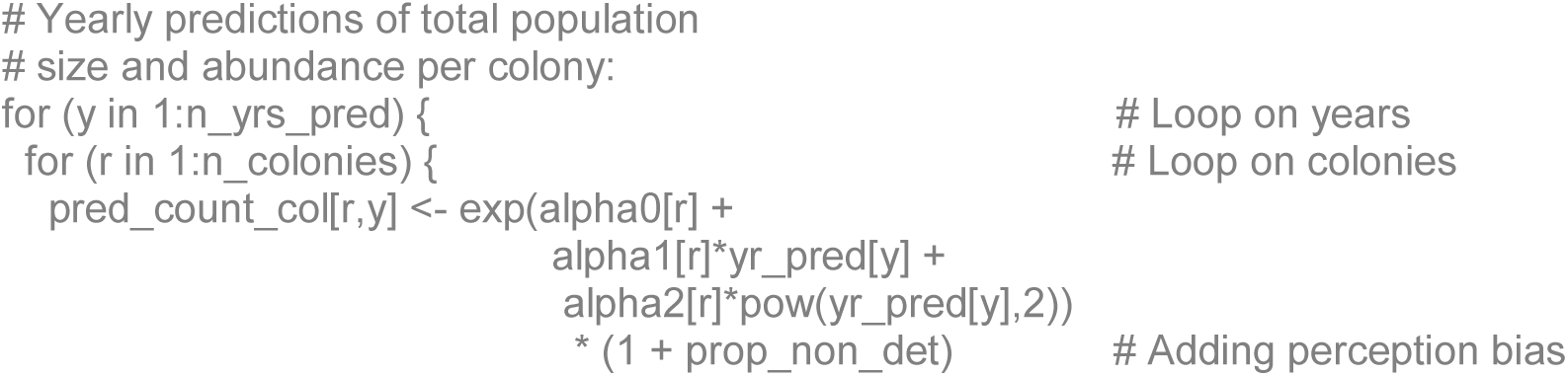

For each year (*y*), a number of adult animals (*Ad*) was estimated at each colony (*r*), using the mean proportion of non-pup animals (*P*_*np*_) defined in Eq. 3:

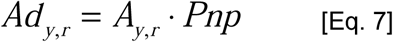

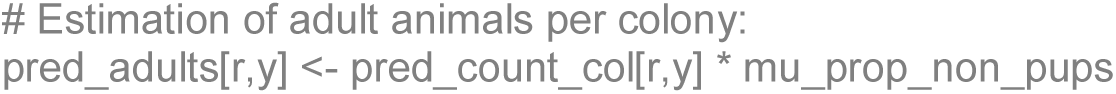

The latter estimate was used to include the correction factor for at-sea animals, not available during the counts, and to obtain the completely corrected annual estimates of abundance (*CA*) for each colony (*r*):

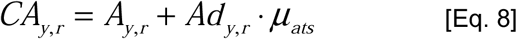

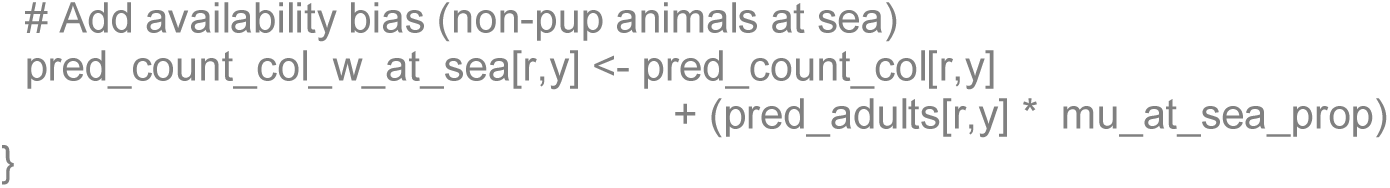

The annual population sizes for the 42 years of the series (*y*) were estimated as the sum of those of the 13 reproductive colonies (*i*), plus the proportion of animals at non-reproductive colonies (*Pnrc*):

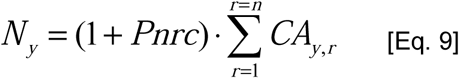

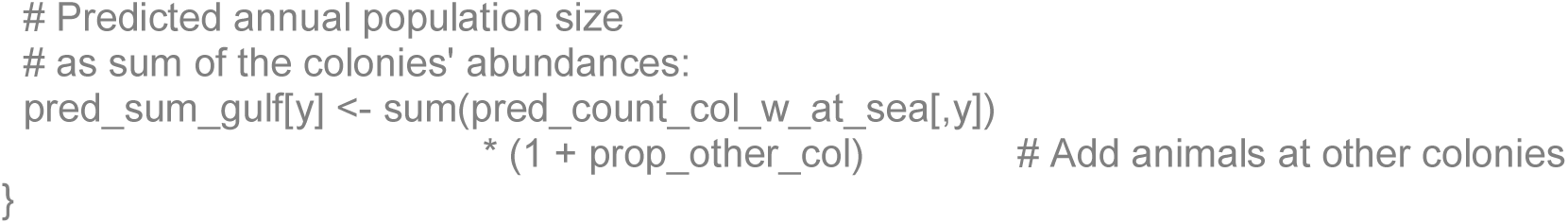

As secondary derived quantities, we estimated the percentage of decadal (*n*) changes in the population size and the main percentage decrease from the highest to the lowest:

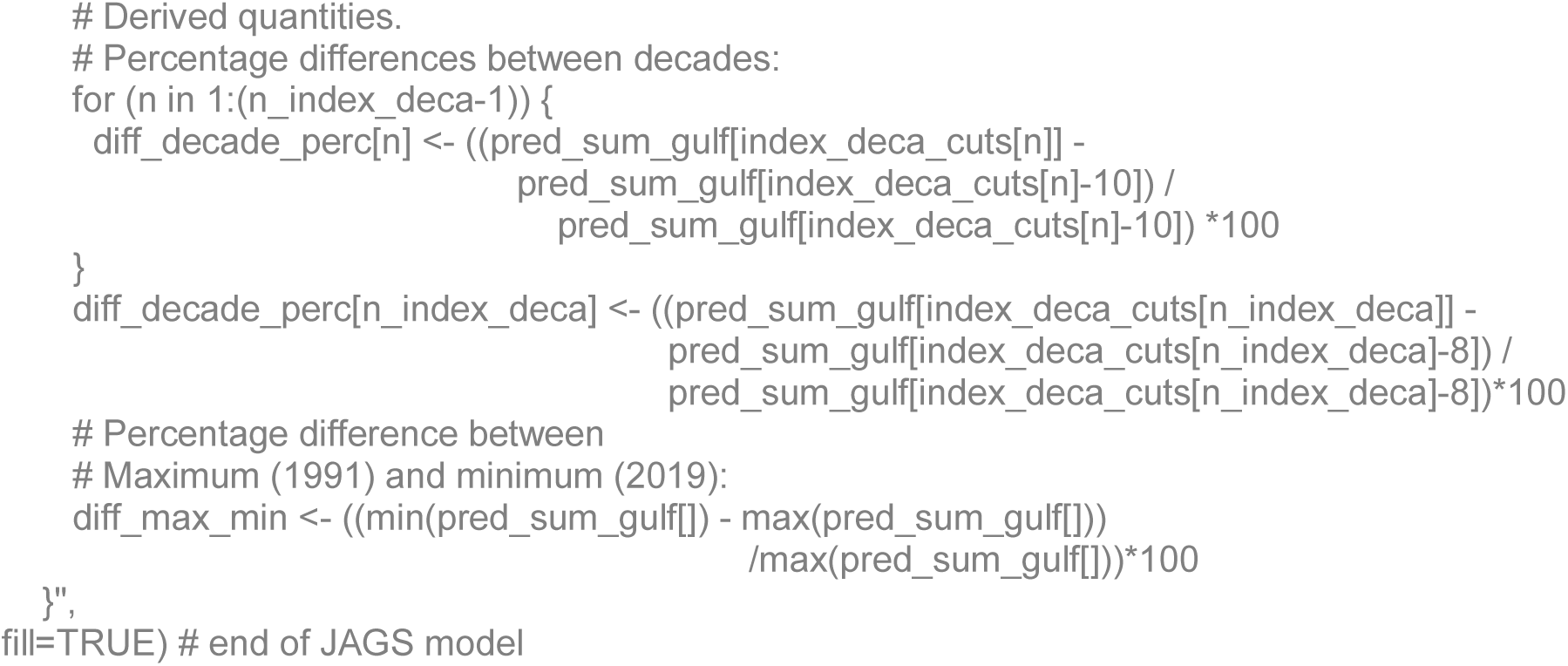

The 100-year analysis of sea surface temperature begins by stating the monthly (*m*) 25-year running means of sea surface temperature (*SST*) to come from a normal likelihood with the 100-year mean (*µ*_*SST*_) and known standard deviations (*σ*_*SST*_; calculated along with the running means). The former was rested to the observations to estimate the anomalies (*SSTa*):

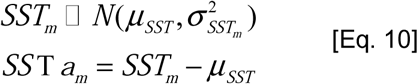

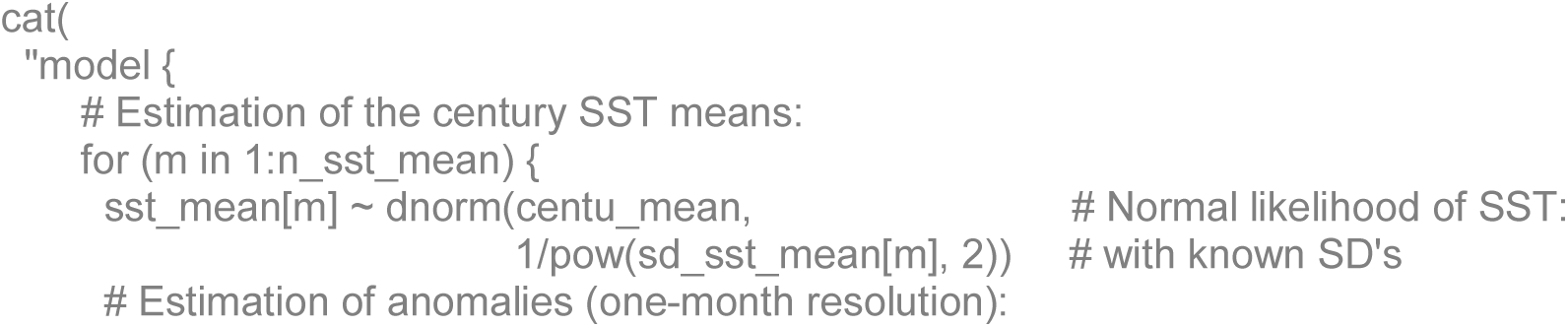

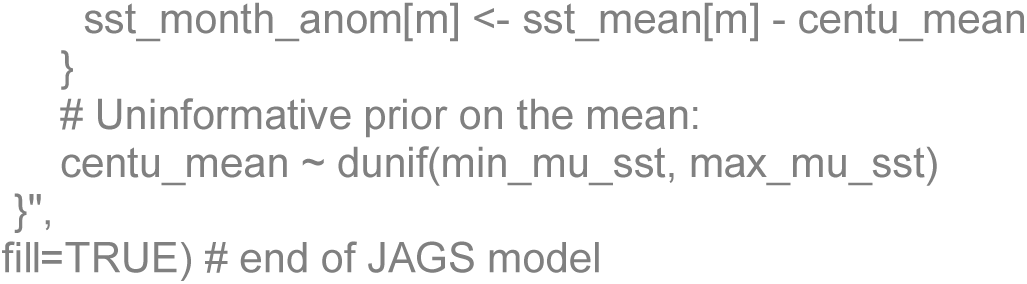

Finally, the ecological model stated that the estimated annual (*y*) populations sizes (*N*) followed a Normal likelihood, whose means were a function of the anomalies (*SSTa)* (only for July estimations). The function with the lowest DIC was a mixture between a parabola and a second-order polynomial with *θ* coefficients, through a mixing parameter (*ω*). The *SSTa* estimations were also stated to come from a Normal likelihood with a mean (*µ*_*SSTa*_) and known standard deviations (*σ*_*SSTa*_):

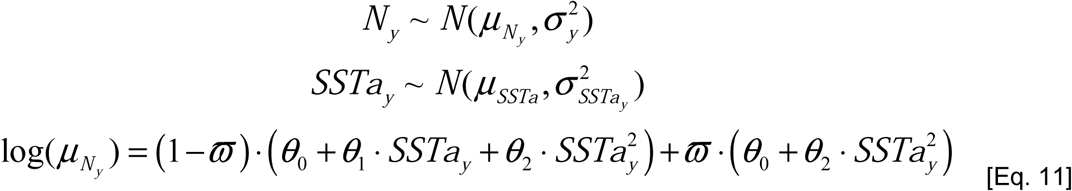

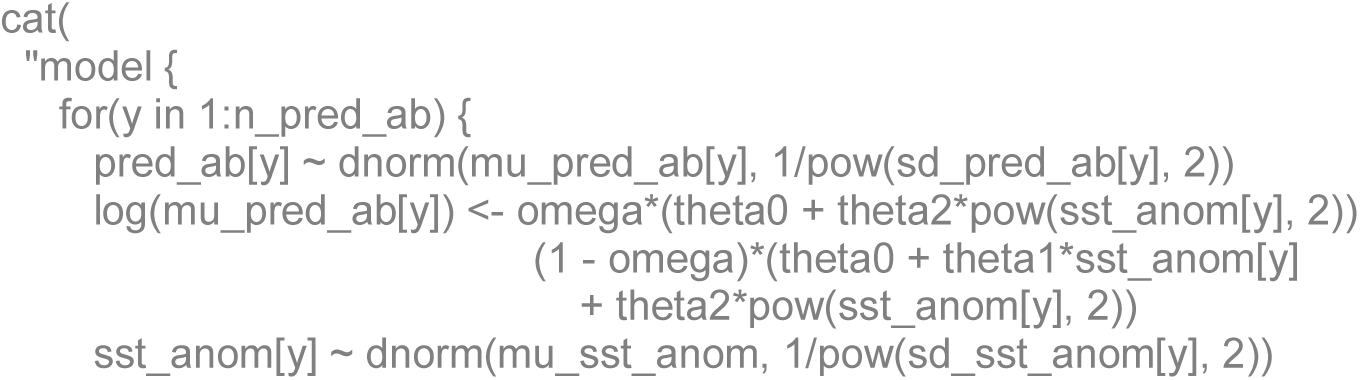

The estimation of the variance explained by this model, the Bayesian R-squared, was based on a vector of errors (i.e. the observed minus the predicted):

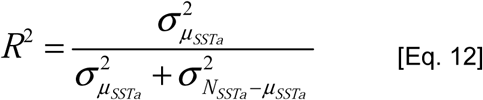

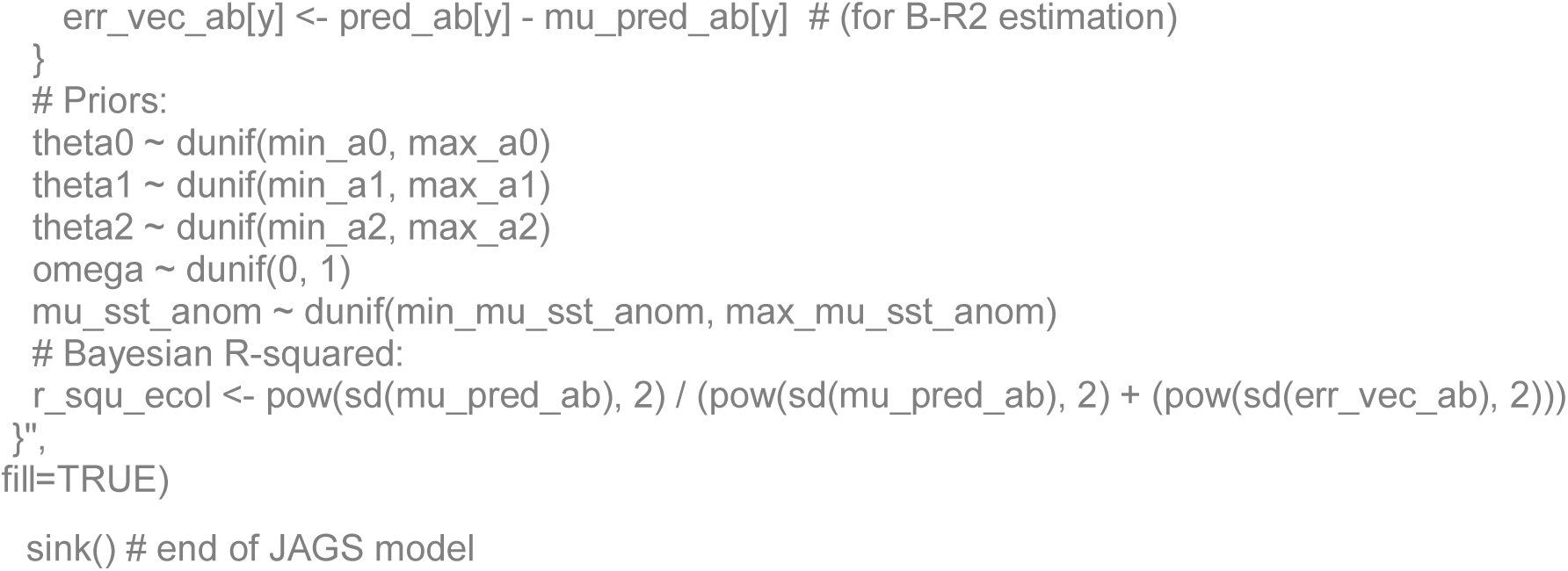

## List of abbreviations

DIC: Deviance Information Criterion
ENSO: El Niño Southern Oscillation
JAGS: Just another Gibbs sampler.
PDO: Pacific Decadal Oscillation
SST: Sea surface temperature

## Declarations

### Ethics approval and consent to participate

This study did not involve any animal manipulation. The animal counts reported for the first time in this study were made during 2016 - 2019 under research permits SGPA/DGVS/00050/16-19 issued by Dirección General de Vida Silvestre - Secretaría de Medio Ambiente y Recursos Naturales, with authorization of Comisión Nacional de Áreas Naturales Protegidas.

### Consent for publication

Not applicable. Most biological data reported in this manuscript has been already published, and was properly referenced. The data reported for the first time belongs to one or more coauthors of this manuscript.

### Availability of data and materials

We assure the technical replication of all the analyses of our study by providing the original biological data (i.e. animal counts) in Supplementary Information Table S1 and File S1, as well as a link within the Methods section granting access to the environmental data (i.e. the SST_ice_; https://coastwatch.pfeg.noaa.gov/erddap/griddap/erdHadISST.html). All the detailed code we built and the equations we used within the analyses, fully replicable in the R language, are given also in the Methods section. Also, all the data necessary for running the three models is provided fully arranged as three R data list objects in the Supplementary Information, File S1. This makes the analyses fully replicable and saves all the programming needed to process and arrange the data.

## Competing interests

The authors declare no competing interests, financial, personal, or other, exist.

## Funding

Funding institutions: CONACYT (Fronteras de la Ciencia; Grant 446; PI: K. Acevedo-Whitehouse), Instituto Politécnico Nacional (SIP grant 20160164; PI: F.R. Elorriaga-Verplancken), and CICESE (Training Scholarship 691102 to K. Adame, and Internal Project 691-113; PI: M.A. Pardo).

## Authors’ contributions

K.A., M.A.P., and F.R.E.V. conceived the study. K.A., F.R.E.V., and K.A.W. were involved in data collection. K.A. processed the sea lion data. M.A.P. carried out the SST_ice_ data processing, all statistical analyses, and figure production. E.B. contributed with the environmental approach. K.A. and M.A.P. wrote the original draft. All authors reviewed and contributed to the final manuscript.

## Acknowledgements

We thank all the personnel who participated in the 2016 Gulf of California survey. E.D. Barton provided useful editorial comments to the original manuscript.

## Notes

https://nimbus.cicese.mx/public.php?service=files&t=de6ad294ec513aabdf9dfe16df544bdd

